# Mathematical Modeling of Field Cancerization through the Lens of Cancer Behavioral Ecology

**DOI:** 10.1101/2023.08.07.552382

**Authors:** Anuraag Bukkuri, Frederick R. Adler

**Affiliations:** Cancer Biology and Evolution Program and Department of Integrated Mathematical Oncology, Moffitt Cancer Center; School of Biological Sciences, University of Utah, Salt Lake City, UT, United States; Department of Mathematics, University of Utah, Salt Lake City, UT, United States; Huntsman Cancer Institute, University of Utah, Salt Lake City, UT, United States

## Abstract

Field cancerization is a process in which a normal tissue is replaced with pre-cancerous but histologically normal tissue. This transformed field can give rise to malignancy and contribute to tumor relapse. In this paper, we create a mathematical model of field cancerization from the perspective of cancer behavioral ecology. In our model, field cancerization arises from a breakdown in signaling integrity and control, and investigate implications for acute wounding, chronic wounding, aging, and therapeutic interventions. We find that restoration of communication networks can lead to cancer regression in the context of acute injury. Conversely, long term loss of controls, such as through chronic wounding or aging, can promote oncogenesis.

These results are paralleled in therapeutic interventions: those that simply target cells in cancerous states may be less effective than those that reestablish signaling integrity. Viewing cancer as a corruption of communication systems rather than as a corruption of individual cells may lead to novel approaches for understanding and treating this disease.

## 1 Introduction

The notion of field cancerization was introduced by Slaughter in 1953 as an explanation for the multitude of oral squamous cell carcinomas, their local recurrences, and the molecular abnormalities observed in hibstologically benign tissue surrounding these cancers [46]. Since then, tumor fields have been discovered in many more organs [6, 12]: head and neck [24, 53], lung [21, 39, 40], vulva and cervix [11, 43], ovaries [9, 38], esophagus [22, 41], skin [15, 18, 30], breast [2, 17, 19], colon [23, 29, 56], stomach [31, 32, 33], gallbladder [51], prostate [14, 52, 57], pancreas [28], and bladder [50].

Also known as field effects or field defects, field cancerization describes carcinogen-induced genetic or epigenetic changes in the epithelium that can give rise to independent malignant lesions, often leading to multifocal tumors [44]. This process occurs in three main steps: First, a carcinogenic insult such as UVA exposure [25], smoke, or alcohol [36] induces genetic or epigenetic aberrations in a cell or population of cells. If these alterations confer a fitness advantage in the existing microenvironment, clonal expansion of the precancerous cells will occur, forming a field. Further alterations transform these cells into malignant, histologically abnormal ones, commonly recognized as cancer. In this environment, removal of cancer cells (e.g., by surgery) does not remove the underlying tumor field, so recurrence is likely.

The field cancerization theory shares many similarities with the multistage model of carcinogenesis [4, 20].

Namely, tumor initiation occurs when the initial cell or cells are transformed, tumor promotion occurs due to the selective expansion of precancerous cells, malignant conversion occurs upon subsequent mutations or epimutations, and tumor progression occurs as malignant cells acquire more aggressive characteristics, promoting regional invasion and eventual distant metastasis. Although the field cancerization theory posits that “cancer does not arise as an isolated cellular phenomenon, but rather as an anaplastic tendency involving many cells at once” [45], field cancerization still puts the focus of cancer at the cellular level.

Here, we adopt a different view. Building on our cancer behavioral ecology work [7], we conceptualize cancer not as an identity problem but as a relational one [47]. Namely, we view cancer as a corruption of communication systems and the potential subsequent selection of “cancerous cells” as a byproduct of this process. Under this framework, cancerous cells are constantly being generated by normal cells, but are quickly eliminated due to controls in the signaling system. When there is a breakdown in communication (e.g., due to a carcinogen, wounding, or aging), these controls are weakened. This selects for cancer cells that clonally expand and create a tumor field from which more aggressive malignancies can develop. In this paper, we create a mathematical model to capture this process and investigate how acute wounding, chronic wounding, aging, and therapy impact tumorigenesis.

## 2 Methodology

### 2.1 ODE Model of Normal, Precancerous, and Malignant Cells

To create a mathematical model of field cancerization, we construct a set of ordinary differential equations (ODEs) to capture the dynamics of three cell populations: normal cells (*N*), precancerous cells (*P*), and malignant cells (*M*). We then simulate the model using a modified Gillespie algorithm, including rare, unidirectional mutations towards malignancy [8]. The ODE model is as follows:

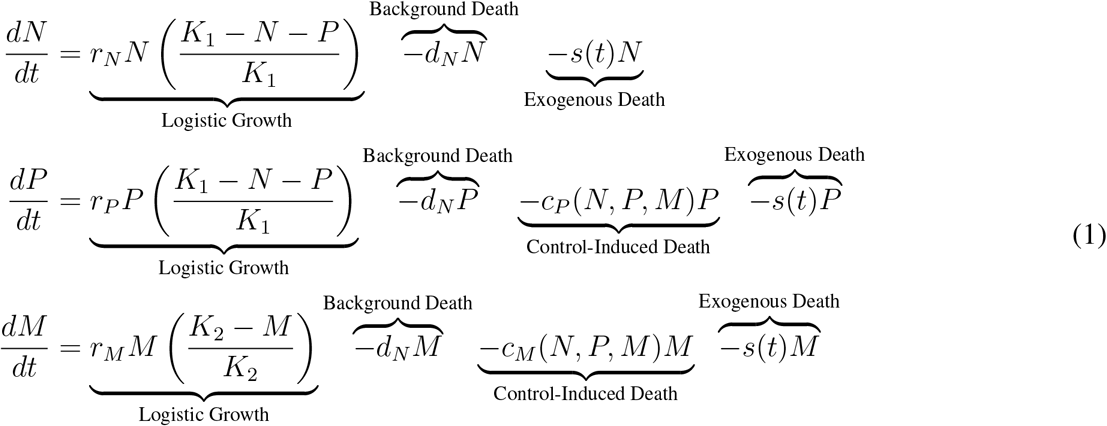

We make the simplifying assumption that all cells grow in a logistic manner, with normal and precancerous cells competing within the epithelium and malignant cells competing only with each other and subject to a larger carrying capacity due to tumor geometry (*K*_2_ *> K*_1_). We let all cells have the same background death rate (*d*_*N*_) and include a death rate for precancerous and malignant cells (*c*_*M*_ *> c*_*P*_) that captures their death due to controls in the signaling system (e.g., immune surveillance). We let

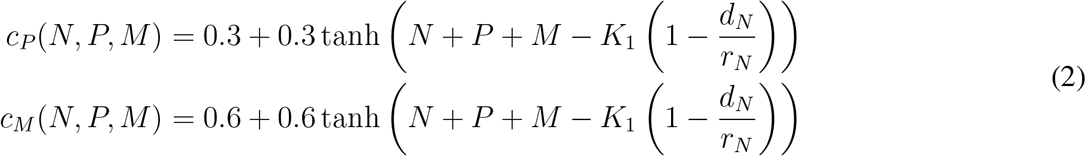

where 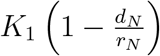 is the number of cells in the tissue at healthy equilibrium. This formulation assumes that immune surveillance, for example, is dampened when the tissue is injured because the need to proliferate and repair the damage outweighs the need to eliminate potential cancer cells. When the tissue returns to homeostasis, the immune system adopts a more discerning phenotype and again removes cancerous cells more effectively. Control-induced death thus varies with the effects of exogenous influences like wounding, aging, or therapy.

Death caused directly by these exogenous factors is captured by *s*(*t*). We assume that the intrinsic growth rate is higher for cells with a more cancerous phenotype: *r*_*M*_ *> r*_*P*_ *> r*_*N*_. The baseline parameter values used in our simulations are given in Table 1.

**Table 1:** Model parameters and values used in simulations.

| Parameter | Interpretation | Value |
| --- | --- | --- |
| $K_1$ | Carrying capacity of normal and precancerous cells | $10^4$ cells |
| $K_2$ | Carrying capacity of malignant cells | $10^5$ cells |
| $r_N$ | Intrinsic growth rate of normal cells | 0.3/day |
| $r_P$ | Intrinsic growth rate of precancerous cells | 0.6/day |
| $r_M$ | Intrinsic growth rate of malignant cells | 0.8/day |
| $d_N$ | Natural death rate | 0.1/day |
| $c_P$ | Control-induced death at homeostasis: pre-cancerous cells | 0.3/day |
| $c_M$ | Control-induced death at homeostasis: malignant cells | 0.6/day |
| $s(t)$ | Environment-induced death | 0/day |
| $\mu_N$ | Mutation rate: normal to precancerous | $10^{-3}$ |
| $\mu_P$ | Mutation rate: precancerous to malignant | $10^{-5}$ |

### 2.2 Stochastic Simulation: Birth-Death-Mutation Process

For simulation, we use a modified Gillespie algorithm, similar to [8]. We first compute birth and death rates at each time step for each cell type from Equation 1. We then compute the total event rate by summing the birth and death rates across cell types. The time to next event is sampled from an exponential probability distribution with a mean of the total event rate. The event type is probabilistically chosen based on the relative contributions of each rate to the total event rate. This event is implemented and the procedure is repeated until a final time is reached and the simulation is stopped. If the event is death, one cell from the respective cell type compartment is eliminated. If the event is birth, we must account for the possibility of mutation. For simplicity, we allow mutations only from normal to precancerous cells (at rate *µ*_*N*_) and from precancerous to malignant cells (at rate *µ*_*P*_). If no mutation occurs, one cell is added to the relevant compartment. If a mutation occurs in a normal cell, one cell is added to the precancerous compartment. If a mutation occurs in a precancerous cell, one cell is added to the malignant compartment. This procedure is summarized in Figure 1.

**Figure 1:**
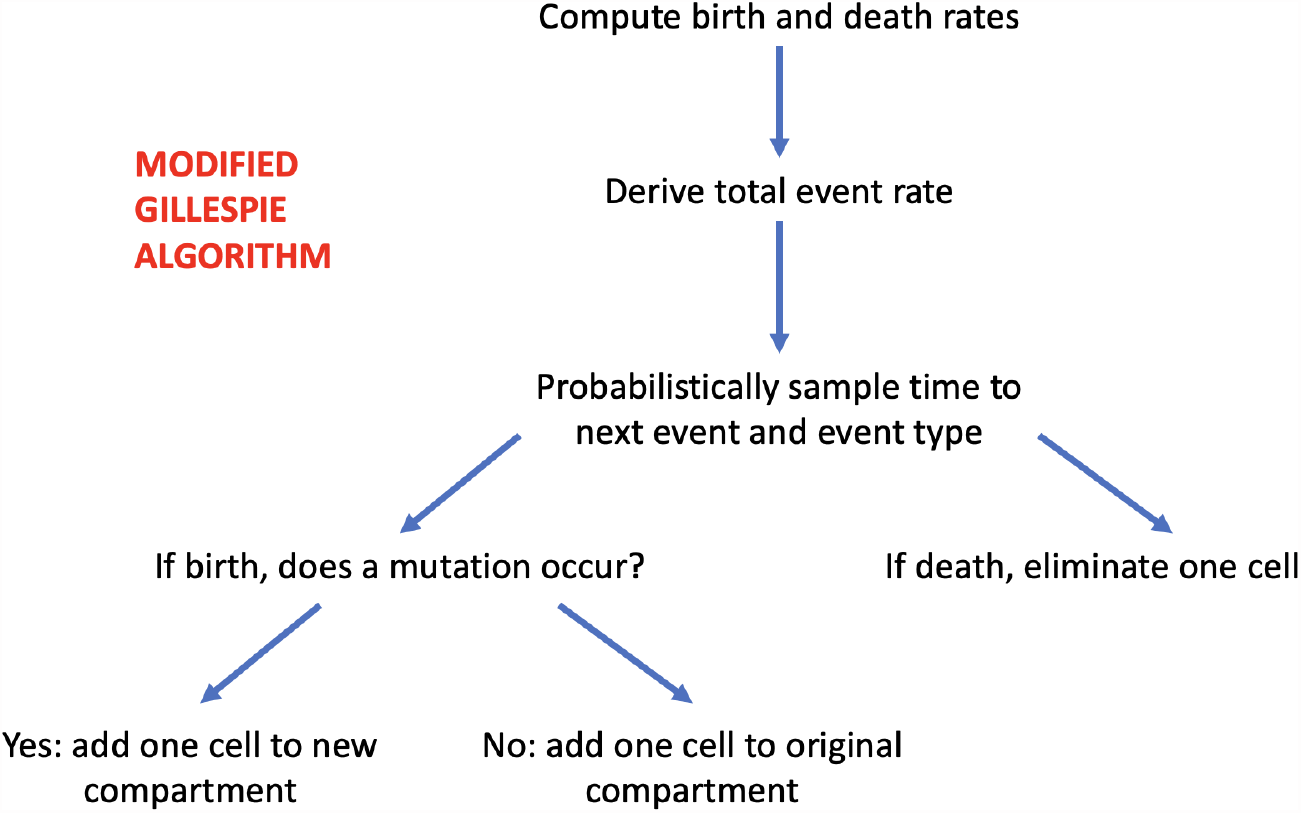
Flowchart depicting the modified Gillespie algorithm used for simulation.

## 3. Results

We simulate the population dynamics of normal, precancerous, and malignant cells under a variety of scenarios. In each case, a set of ten simulations are performed as follows: We initialize the population at homeostatic equilibrium (6667 normal cells) and run the simulation for 750 days. We test whether the system transiently or permanently switches to a cancerous state in four scenarios: healthy tissue homeostasis, acute wounding, chronic wounding, and aging. Finally, we show how restoration of controls is more effective than removal of cancerous cells for cancer elimination.

### 3.1 Control: Healthy Case

In the healthy case, the tissue is at homeostasis. The communication network is robust. The high penalty on cells in a precancerous or malignant state should maintain a population composed primarily of normal cells. We implement this scenario in Fig. 2 by using the default parameter values given in Table 1. As expected, normal cells dominate the population at all times. Although cancerous cells occasionally arise, high control-induced death rates lead to their rapid elimination.

**Figure 2:**
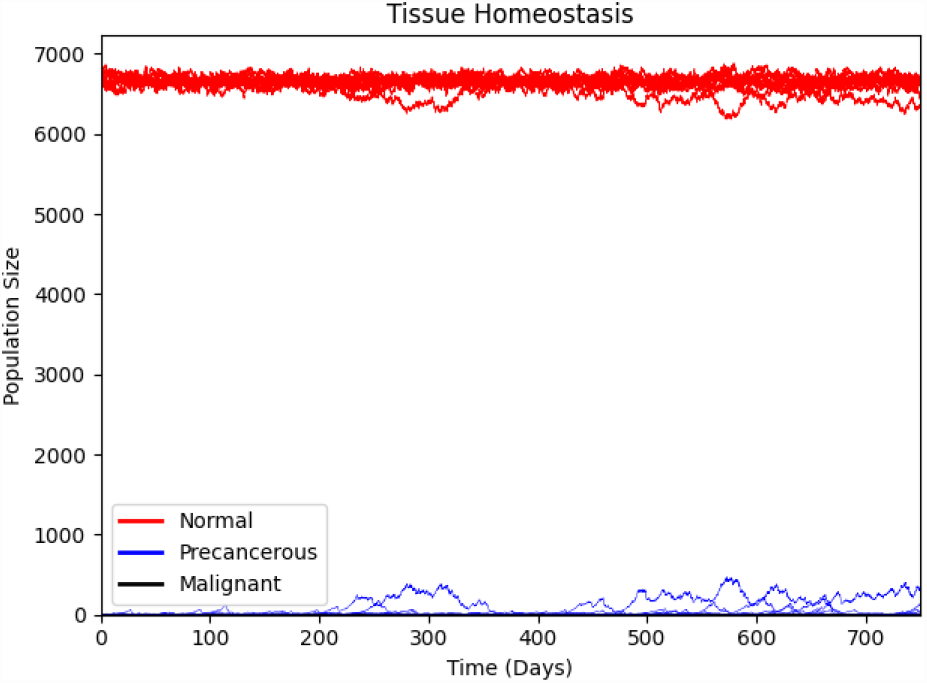
Healthy control. Red, blue, and black lines represent normal, precancerous, and malignant cells respectively. Each line corresponds to one, out of a total of ten, replicate simulation runs. Despite high cell division rates, precancerous and malignant cells exist at very low frequencies in the population due to high control-induced death rates. The tissue is composed of nearly all normal cells at all times.

### 3.2 Acute Wounding

With acute wounding, the tissue suffers damage due to a traumatic event which leads to the death of most its cells. In addition, cell communication networks and morphogenetic fields are disrupted and immune surveillance is dampened, leading to the transient loss of controls. As the tissue is repaired, controls are gradually reinstated. This differs from chronic wounding (Section 3.3) in two ways: the stressor only occurs once (allowing for tissue recovery and restoration of signaling integrity) and tissue damage due to the wound is severe. To simulate this, we induce the traumatic event at day 100 by setting *s*(*t*) = 1 for *t∈* [100, 101]. This kills roughly 37% of the cells in the tissue.

Before the wound, the healthy tissue is comprised almost entirely of normal cells. Once the wound occurs, the population size drops dramatically due to rapid cell death. Signaling integrity is disturbed and controls are lost, leading to the expansion of cancerous cells. However, as the communication networks are restored and controls are put back in place, cells revert to their original frequencies. Restoring signaling networks is thus a potent way to promote cancer regression.

### 3.3 Chronic Wounding

Chronic wounding continually damages the tissue due to a chronic stressor such as UV radiation, tobacco, diabetes, or ulcers. This leads to excessive inflammation and disruption of signaling integrity in the tissue, ultimately hindering tissue repair and promoting tumorigenesis. This builds on the long-standing description of cancer as a “wound that never heals” [10, 13, 16, 26, 55]. Unlike the tightly regulated process of recovery from acute wounding, tissue injury is never resolved and controls are not reinstated. Thus, the self-limiting hyperproliferative behavior during epithelial regeneration is left unchecked, allowing growth of cancer [49]. To simulate this phenomenon, we assume that chronic wounding begins at day 100 and let *s*(*t >* 100) = 0.15 to capture continual tissue damage (Fig. 4).

**Figure 3:**
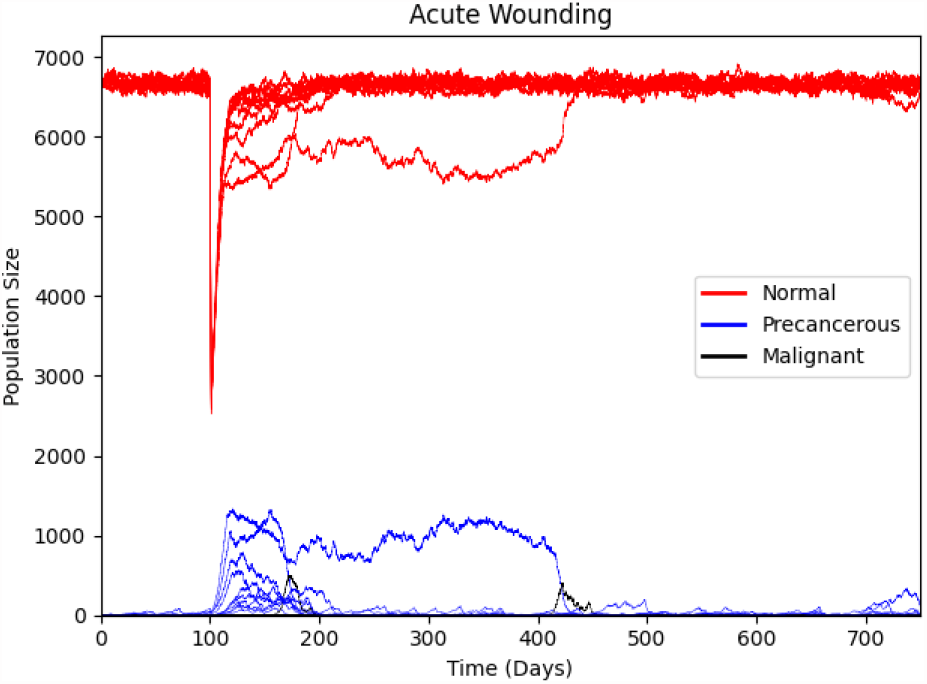
Acute wounding. Red, blue, and black lines represent normal, precancerous, and malignant cells respectively. Each line corresponds to one, out of a total of ten, replicate simulation runs. A traumatic event leads to rapid cell death. This leads to a loss of controls and allows for the expansion of precancerous and malignant cells. As signaling integrity is gradually restored, these cancerous cells are eliminated and the tissue returns to healthy homeostasis.

**Figure 4:**
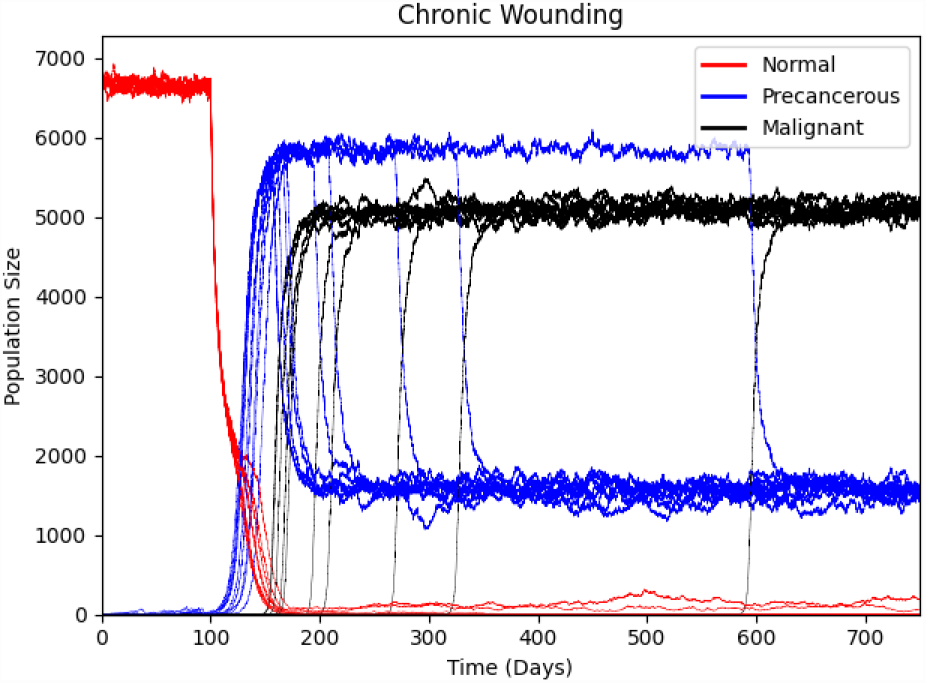
Chronic wounding. Red, blue, and black lines represent normal, precancerous, and malignant cells respectively. Each line corresponds to one, out of a total of ten, replicate simulation runs. Chronic wounding causes modest cell death, high cell turnover, and a corruption of communication networks. This paves the path for precancerous and malignant cells to invade the tissue.

After establishing healthy homeostasis during the first 100 days, chronic wounding reduces the population size, thereby disrupting signaling systems and weakening controls. This allows for the expansion of precancerous and ultimately malignant cells, that eventually overtake the tissue (Fig. 4).

### 3.4 Aging

Aging can be viewed as a gradual loss of controls and signaling integrity due to sarcopenia, tissue structural atrophy, and a weakening of immunosurveillance abilities [5, 34, 35, 54]. To simulate this process, we let control-induced death be a decreasing function of time: *c*_*P*_ (*N, P, M, t*) := *c*_*P*_ (*N, P, M*) exp(*−*0.01*t*) and *c*_*M*_ (*t*) := *c*_*M*_ (*N, P, M*) exp(*−*0.01*t*) (Fig. 5). The start of the simulation parallels that of healthy homeostasis: the communication system has not been disturbed and cancerous cells are unable to invade the population. Eventually, controls weaken to the extent that precancerous and malignant cells take over the population. By the end of the simulation period, no healthy cells remain. Note that the total population size far exceeds that in our other simulations. This is due to the gradual decimation of controls: By the end of the simulation, control-induced death is negligible. This allows the malignant cells to grow unencumbered by the effects of the immune system, only limited in size by their carrying capacity.

**Figure 5:**
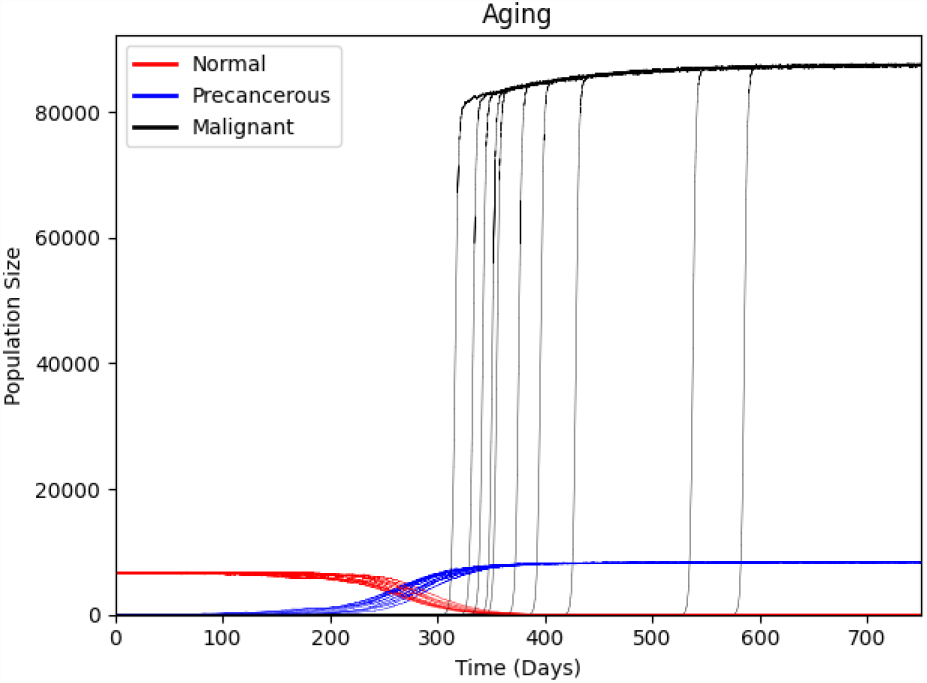
Aging. Red, blue, and black lines represent normal, precancerous, and malignant cells respectively. Each line corresponds to one, out of a total of ten, replicate simulation runs. As controls gradually weaken over time, precancerous and malignant cells expand to dominate the population.

### 3.5 Therapeutic Interventions

With this model of cancer initiation and progression through the lens of field cancerization and cancer behavioral ecology, we now consider implications for therapy. We investigate two therapeutic approaches. The first eliminates cells in the precancerous and malignant states via surgery. The second restores controls and signaling network integrity such as through immunotherapy [1, 37, 48]. In both cases, we will induce cancerization by setting *c*_*P*_ = *c*_*M*_ = 0. Therapeutic interventions are introduced when the cancerous population size exceeds the normal population size and the simulation is run until day 750.

First, we consider the traditional approach of eliminating cells in the cancer state. Focusing on surgical interventions, we make the extreme assumption that radical surgery removes all malignant and precancerous cells, while leaving all normal cells intact. To implement surgery, we set *P* = *M* = 0 (Fig. 6). Surgery temporarily removes the cancer, but recurrence is inevitable because the lack of controls reinstates a cancer field, giving rise to malignancy.

**Figure 6:**
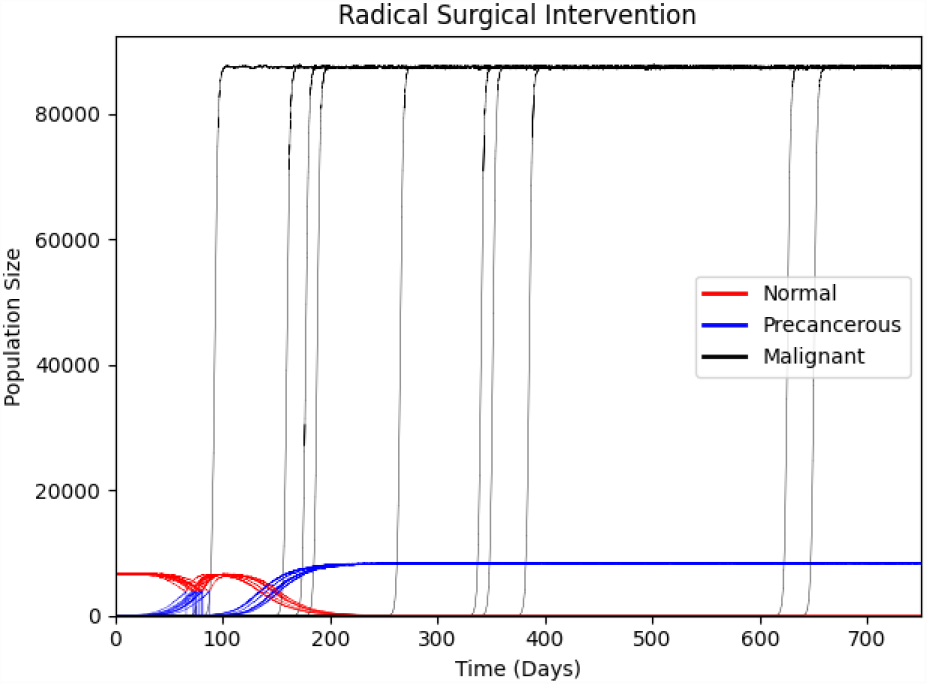
Impact of radical surgery. Red, blue, and black lines represent normal, precancerous, and malignant cells respectively. Each line corresponds to one, out of a total of ten, replicate simulation runs. Total elimination of precancerous and malignant cells cannot prevent tumor recurrence.

We hypothesize that therapeutic interventions that focus on reestablishing signaling integrity (e.g., via CAR-T therapy [48], macrophage polarization [3], tumor microenvironment modification [42], or reinstating self-regulatory feedback loops on growth factor and cytokine production [13, 27, 49]) rather than directly killing cells in a cancerous state will be more effective for tumor elimination. To implement this strategy, we restore our controls to *c*_*P*_ = 0.3 and *c*_*M*_ = 0.6 when the cancerous population size surpasses the normal population size (Fig. 7). As expected, restoration of controls leads to high control-induced death rates in cancerous cells, leading to their elimination and preventing tumor recurrence.

**Figure 7:**
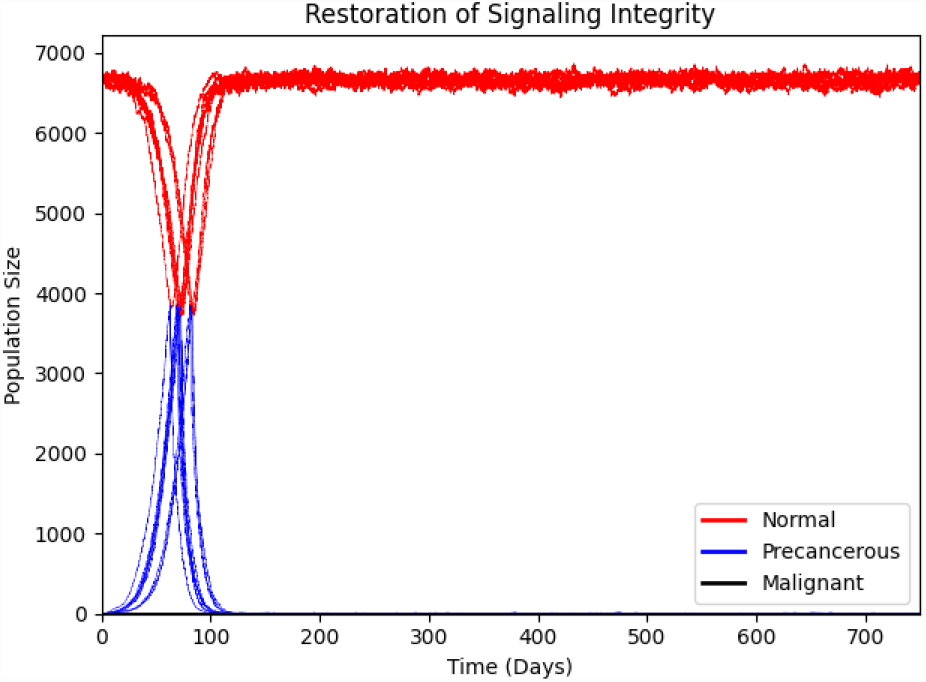
Impact of restoring signaling integrity. Red, blue, and black lines represent normal, precancerous, and malignant cells respectively. Each line corresponds to one, out of a total of ten, replicate simulation runs. Fortification of communication networks leads to rapid tumor regression.

## 4 Conclusion

Current cancer therapeutic efforts focus on effective ways of killing cells in a cancerous state. Although this approach may be effective for elimination in the short term, it often allows recurrence. Here, we approached oncogenesis through a field cancerization lens and view oncogenesis as a three-step process: initial transformation via genetic or epigenetic aberrations, clonal expansion of precancerous clones to form a cancer field, and further transformations towards malignancy. Rather than adopting the traditional view that cancer is a disease of the cells, we take a cancer behavioral ecology approach and view cancer as a corruption of signaling systems. In this view, cancerous cells are continuously being generated within a tissue, but are quickly eliminated due to robust cell communication networks. When these networks are disrupted, “cancer cells” escape control and thrive.

In this paper, we create a mathematical model to investigate the consequences of this world view for disruptions of signaling integrity by acute wounding, chronic wounding, and aging. Long term loss of controls does indeed lead to the expansion of cancerous cells. We also examine the efficacy of two classes of therapeutic interventions: traditional strategies that eliminate cancer cells and strategies that fortify the signaling networks themselves, finding that restoration of signaling integrity is the key to prevent relapse.

Future work will expand on this model in several directions. We plan to construct an explicit consumer-resource model to allow the growth rate of normal, precancerous, and malignant cells to depend on growth factors in the microenvironment, a key mechanism in wounding and inflammation. These models will include explicit mechanisms for how cancer cells hijack, corrupt, and co-opt communication networks in tissues and for tumor-immune and epithelial-stroma interactions.

## 5 Data Availability

Codes associated with the plots produced in this paper can be found at https://github.com/abukkuri/FieldCancer.

